# A versatile, multi-laser twin-microscope system for light-sheet imaging

**DOI:** 10.1101/801688

**Authors:** Kevin Keomanee-Dizon, Scott E. Fraser, Thai V. Truong

**Affiliations:** Translational Imaging Center, Dornsife College of Letters, Arts and Sciences, and Viterbi School of Engineering, University of Southern California, Los Angeles, CA 90089, USA

## Abstract

Light-sheet microscopy offers faster imaging and reduced phototoxicity in comparison to conventional point-scanning microscopy, making it a preferred technique for imaging biological dynamics for durations of hours or days. Such extended imaging sessions pose a challenge, as it reduces the number of specimens that can be imaged in a given day. Here we present an instrument, the flex-SPIM, that combines two independently controlled light-sheet microscope-twins, built so that they can share an ultrafast near-infrared laser and a bank of continuous-wave visible lasers, increasing throughput and decreasing cost. To permit a wide variety of specimens to be imaged, each microscope-twin provides flexible imaging parameters, including (i) operation in one-photon and/or two-photon excitation modes, (ii) delivery of one to three light-sheets *via* a trio of orthogonal excitation arms, (iii) sub-micron to micron imaging resolution, (iv) multicolor compatibility, and (v) upright and/or inverted detection geometry. We offer a detailed description of the flex-SPIM design to aid instrument builders who wish to construct and use similar systems. We demonstrate the instrument’s versatility for biological investigation by performing fast imaging of the beating heart in an intact zebrafish embryo, deep imaging of thick patient-derived tumor organoids, and gentle whole-brain imaging of neural activity in behaving larval zebrafish.

## I. INTRODUCTION

Most of what we recognize as the phenomena of life are not properties of stationary structures, but rather emerge from dynamic interactions among many elements over time. Modern optical microscopy methods offer an efficient means for non-invasive, high-resolution observation of many of life’s most fascinating phenomena. The difficulty is that light imaging involves unavoidable tradeoffs between spatial resolution, acquisition speed, field-of-view, penetration depth, and the limited photon budget from the sample.^1^ Considerations of photon budget are crucial to biological imaging, as there are a finite number of photons that a given fluorophore can emit before it is bleached, and high light doses on the specimen can lead to photo-induced toxicity. It is thus critical that the excitation light be used as efficiently as possible and the emitted photons be collected as efficiently as possible, while striking the optimal imaging compromises for whatever the application demands.

Over the past decade, there have been a series of important developments in light-sheet microscopy, a century-old technique^2^ (also known as selective-plane illumination microscopy; SPIM).^3^ SPIM decouples the illumination and detection paths by using separate optics to excite and detect fluorescence: a cylindrical illumination lens is used to project a static, thin, two-dimensional (2D) sheet of light coincident to the focal plane of a detection objective lens. In contrast to confocal laser scanning microscopy and other point-scanning techniques that acquire volumetric information one voxel at a time, light-sheet excitation permits an entire 2D plane of fluorophores to be excited and detected with high signal-to-noise ratio, high imaging speed, and minimal light exposure to the sample. Developments in light-sheet microscopy techniques^4–11^ have led to cutting-edge applications across a range of fields,^4–7,10,12,13^ from developmental biology^5,14^ to neuroscience.^10,12,15–17^ Each of these modifications of light-sheet imaging has required their own tradeoffs in performance, complexity and expense of the microscope optics, and expenditure of the photon budget. Comprehensive reviews of light-sheet development can be found elsewhere (*e.g.* see Ref 5)—we highlight below several key developments that address the experimental demands which motivate the development of our instrument.

A key development was to create light-sheets by dynamically scanning a focused Gaussian beam, generated *via* a low numerical aperture (NA) lens, across the plane (DSLM; Fig. 1).^18^ This scanned Gaussian-beam light-sheet approach provides better spatial illumination uniformity, higher light throughput, and more precise spatial control over the selected plane of interest compared to a static 2D light-sheet, at the cost of replacing the simple cylindrical lens with the expense and complexity of a scanning galvanometer (galvo) mirror, and associated optics and electronics. This task has been simplified by the commercial availability of integrated electro-optical galvo scanning modules.

**FIG. 1.**
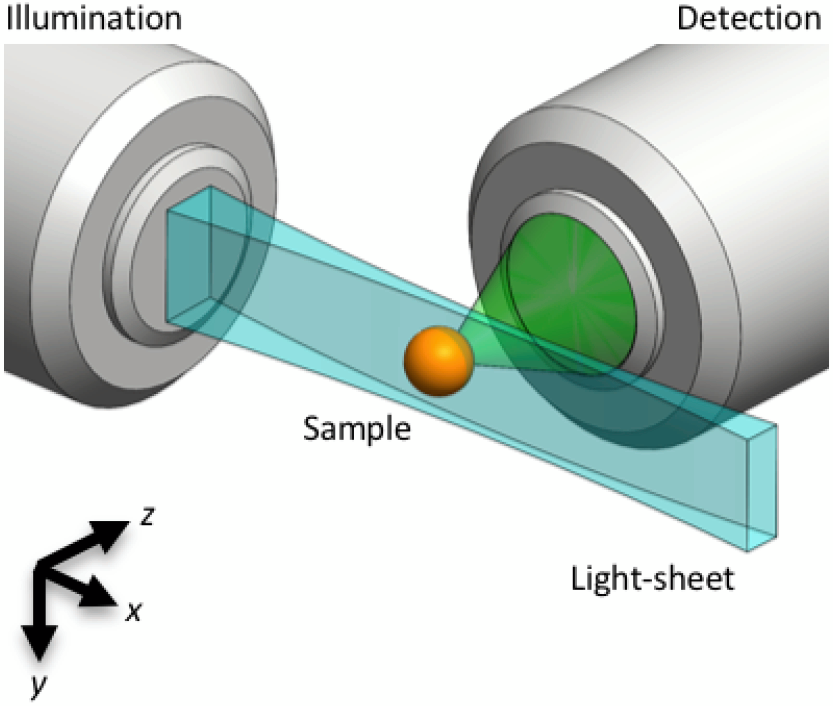
Light-sheet microscopy principle. A light-sheet (blue) can be created by dynamically scanning, along the *y* direction, a focused Gaussian beam which propagates in the *x* direction. The focusing is achieved *via* a low numerical aperture illumination lens. The illuminated (*x-y*) plane is detected (green) by an orthogonally positioned wide-field microscope. Axial (*x-y*) sections are captured either by scanning the sample (orange) through the stationary focal plane, or by scanning the light-sheet and detection focal plane through the stationary sample.

The penetration depth of DSLM was improved in two-photon light-sheet microscopy (2P-SPIM) using nonlinear excitation.^19^ 2P-SPIM has proven successful in relatively thick or optically dense samples, imaging up to 2-fold deeper than DLSM (1P-SPIM), and more than 10-fold faster and 100-fold lower peak intensity than conventional 2P point-scanning microscopy. In addition, the near-infrared (NIR) light used for 2P excitation is invisible to many animals, which avoids unintended visual stimulation.^20^ Both 1P- and 2P-SPIM imaging can be conveniently carried out on the same setup, since both use a low NA illumination lens to generate an axially-extended Gaussian focus.^19,21^

A major expense of SPIM imaging set-ups is the laser source used for excitation, especially given that multiple lasers are used for multicolor imaging. For 2P-SPIM the ultrafast laser can more than double the equipment cost, which has limited its adoption, despite its superior performance in optically challenging samples. Even for well-funded laboratories, laser sources tied to a single microscope are not cost-effective. Since there is an upper limit to the amount of laser power that can be delivered to any specimen without perturbing it, most implementations waste well more than half of the total laser power available.

Another practical challenge comes from the need to continuously image a large number of samples in order to obtain statistically significant results. While the low photodamage of SPIM allows biological processes to be imaged for a duration of several hours or days, a tradeoff exists between such prolonged imaging sessions and the number of samples that can be imaged in a given day using a single instrument.

Here we describe the flex-SPIM, which combines two independently controlled light-sheet microscope-twins that share an ultrafast NIR laser and a bank of continuous-wave (CW) visible lasers. This permits two specimens to be imaged simultaneously for far less than the cost of two multi-laser microscopes. Each microscope-twin has built-in modularity for tailoring its use on diverse samples and scientific questions. In the following section we describe the flex-SPIM design in detail, for those who wish to construct and use a similar instrument. We test the performance against our design objectives by imaging three challenging samples: the beating larval zebrafish heart, patient-derived tumor organoids, and whole-brain neural dynamics in behaving zebrafish. This demonstrates the ease of adapting the flex-SPIM for application-specific light-sheet imaging.

## II. INSTRUMENT DESIGN, INTEGRATION, & CONFIGURATION

The flex-SPIM draws on lessons learned from proof-of-principle studies^19,21^ with a new imaging technology to meaningful scientific results;^22–24^ several years of interactions with end users at advanced imaging centers (at the California Institute of Technology and the University of Southern California), the 2P-SPIM inventors, and other instrument builders; and thus integrates the following combination of improvements:

- Two independent microscope-twins share the same multi-laser source (Fig. 2), dramatically reducing instrument cost—a more than 30% savings.
- The twin architecture doubles the 1P- and 2P-SPIM imaging throughput, and increases the variety of specimens or imaging modes, when compared to a single microscope.
- The system is “flexible” by design, with an opto-mechanical configuration that is both open and modular, providing a straight-forward path to instrument evolution and customization for different samples and applications. Three orthogonal illumination arms offer easy matching to different specimens, enhancing illumination uniformity or increasing optical coverage for larger and more opaque samples.^25^ Switching from high lateral spatial resolution (sub-micron) to a lower spatial resolution (~ micron) with a larger field-of-view requires only a simple adjustment to the detection subsystem. Each twin can be configured in upright and/or inverted detection geometries to accommodate a diversity of specimens (Fig. 2).

**FIG. 2.**
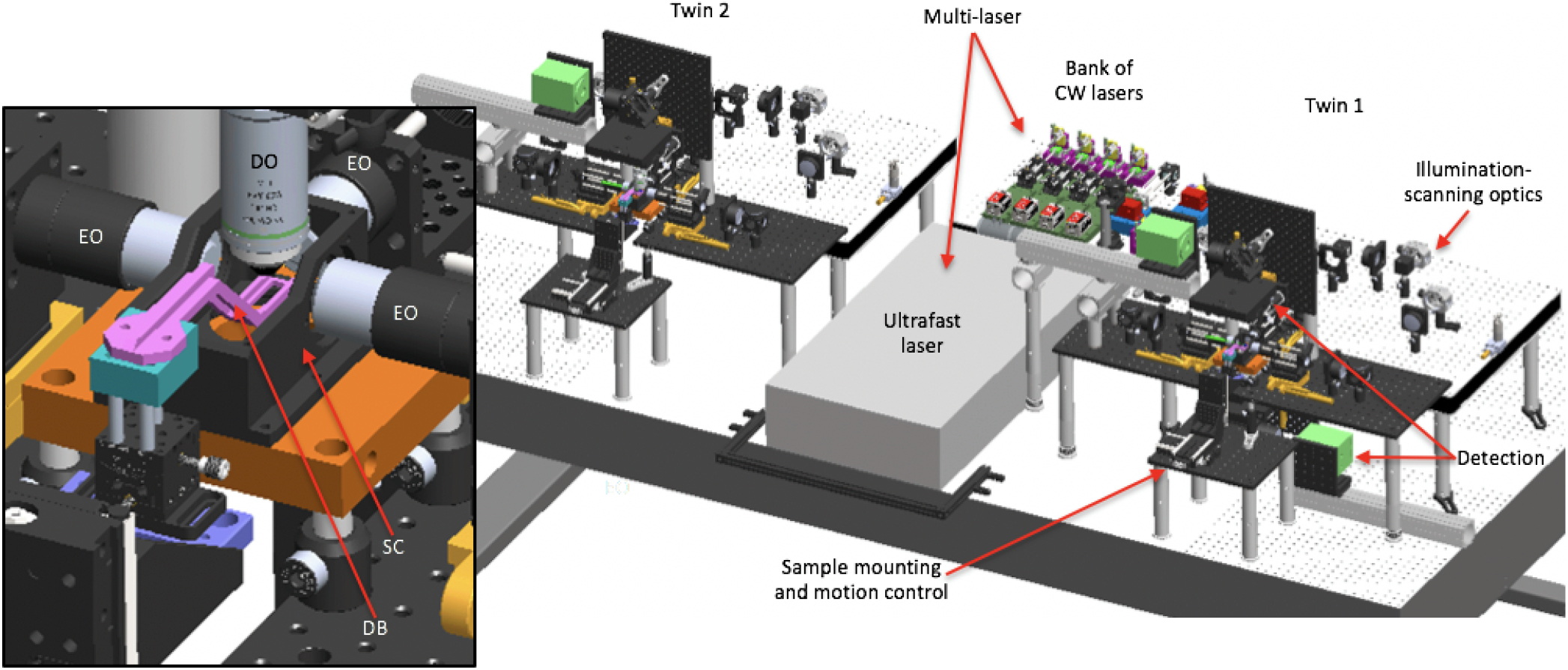
3D opto-mechanical model of the twin-microscope system mounted on a 5 foot × 10 foot, anti-vibration optical table. Model shows the multi-laser subsystem shared between microscope-twin-1 (right) and microscope-twin-2 (left). Twin-1 has the four functional subsystems labeled and features an implementation of both upright and inverted detection. Brief descriptions of each functional subsystem are provided in Table I. The inset shows a detailed view of the sample chamber (SC); the dive bar (DB) used to hold the sample; three excitation objectives (EO) to deliver excitation light-sheets to the sample; and the detection objective (DO) to collect emitted fluorescence from the sample.

The flex-SPIM consists of four functional subsystems (Fig. 2) and two modules (Table I), and sits on a 5 foot × 10 foot, anti-vibration optical table (Fig. S1). The schematic diagram of the integrated illumination paths is shown in Fig. 3, and the corresponding 3D opto-mechanical model is shown in Fig. 2 and available upon request. Whenever possible, commercially available hardware components are used; however, both basic machining of off-the-shelf parts and fabrication of custom components are required (Table SI). Most standard optical elements are mounted in Thorlabs 30 mm or 60 mm cage components. Beam steering mirrors shared by both the ultrafast and CW lasers (illumination-scanning optics) have protected silver coatings, whereas those used by the ultrafast or CW lasers alone have broadband dielectric coatings.

**TABLE I.** flex-SPIM functional subsystems and modules. CW: continuous-wave; polarization beamsplitting optics: half-wave plate and polarizing beamsplitter; AOTF: acousto-optic tunable filter; galvo: galvonometer; LED: light-emitting diode

| Subsystem/module | Description | Main components<sup>a</sup> |
| --- | --- | --- |
| Multi-laser | Laser modulation and beam routing | CW lasers; beam expander; dichroic mirrors; polarization beamsplitting optics; AOTFs; mirrors<br>Ultrafast laser; polarization beamsplitting optics; beam expander; Pockels cells; mirrors |
| Illumination-scanning optics | Generating the light-sheet | Scanning galvo mirrors and optics; mirrors; objectives |
| Detection | Image capture | Scientific camera; tube lens; filter wheel; filters; objective |
| Sample mounting and motion control | Holding and imaging the 3D sample | Custom sample chamber/holder; motion stages; piezoelectric z-stage |
| Instrument control | Control and timing of components | Acquisition computer; National Instruments PXI; system mainframe and scaling amplifiers |
| Auxiliary | Illumination masks and view-finding | Camera; LED; filter; aluminum masks |

<sup>a</sup> The main parts for each subsystem and module used to build the flex-SPIM are listed in Table SI.

**FIG. 3.**
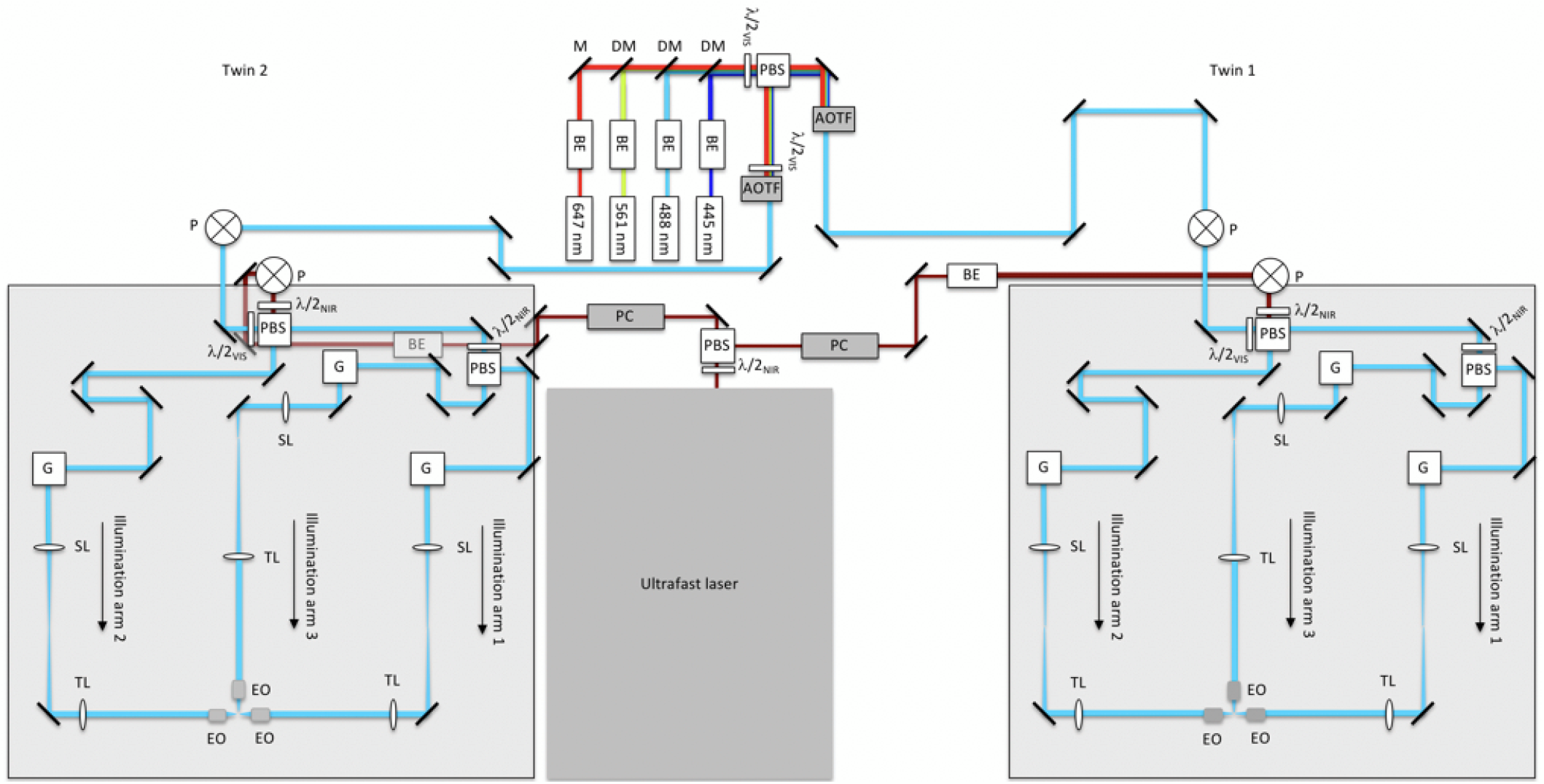
Schematic diagram of multi-laser and illumination-scanning optics subsystems of the instrument. Visible light from the continuous-wave laser bank is fed into microscope-twin 1 (right) and microscope-twin 2 (left) *via* polarization beamsplitting optics [consisting of a polarizing beamsplitter (PBS) and half-wave plate]. Acousto-optic tunable filters (AOTFs) are used to select the visible wavelengths and adjust the power independently for each twin. The near-infrared (NIR) light from the ultrafast laser is routed similarly using Pockels cells (PCs) to adjust the NIR power independently for each twin. Both the visible and NIR beams are each raised onto 24 inch × 36 inch optical breadboards by periscopes (P). Polarization beamsplitting optics are used to both combine the visible and NIR beams, and to split the combined beam into two paths (illumination arms 1 and 2). Illumination arm 1 is further split into two paths through polarization beamsplitting optics, creating a total of 3 illumination arms. Each illumination arm directs light to the sample through the excitation objectives (EO). BE: beam expander; M: mirror; DM: dichroic mirror; *λ*/2: half-wave plate, where the subscripts VIS and NIR refer to the visible and near-infrared wavelengths, respectively; G: 2D scanning galvo mirrors; SL: scan lens; TL: tube lens. BE in the NIR twin-2 path appears gray because it is underneath the optical breadboard.

### A. Multi-laser subsystem

CW visible light used for single- and multicolor imaging *via* linear excitation is provided by a bank of CW lasers (445 nm, 488 nm, 561 nm, and 647 nm), collimated and expanded to a 1/e^2^ diameter of 1.5 mm, and combined into a co-linear beam using broadband and dichroic mirrors (see Table SI).^26^ The combined beam is then split into two paths of equal length and power through polarization beamsplitting optics (consisting of a half-wave plate mounted in a rotation mount and polarizing beamsplitter), delivering light to each microscope-twin. Each light path passes through acousto-optic tunable filters (AOTFs), which are used to select the wave-lengths and adjust the power independently for each twin. The AOTFs (Table SI) require the input laser beam polarization to be linear orthogonal to the baseplate (s-polarization), to maximize the diffraction efficiency and ensure chromatic co-linearity of the modulated beam. Alternatively, the AOTF input beam can be p-polarized if the crystal output face is used as the “input” face due to the Helmholtz reciprocity principle.^27^ Because of the upstream polarizing beamsplitter used, the beams are p-polarized for the twin-1 path and spolarized for the twin-2 path. As such, the AOTF in the twin-1 path is mounted so that the p-polarized beam enters the AOTF through its output face; the AOTF in the twin-2 path is mounted conventionally. A half-wave plate can be placed in front of the AOTF to fine-tune the polarization direction of the beam entering the AOTF, and thereby maximize the diffraction efficiency by the AOTF when more excitation energy is required at the sample, as we have implemented in the twin-2 path (Fig. 3). Note that alternatively, similar performance could be achieved by placing a half-wave plate in front of the AOTF in the twin-1 path to rotate the beam’s polarization so that it enters the AOTF conventionally.

The tunable NIR ultrafast mode-locked laser used for single- and multicolor imaging *via* nonlinear excitation is split into two paths of equal length and power through polarization beamsplitting optics, delivering NIR light to both microscope-twins. Note that the linear polarizations of the beams for the twin-1 path and the twin-2 path are orthogonal to each other. Each path passes through a Pockels cell to control the power independently for each twin (Fig. 3). Each Pockels cell is rotated to match its input polarization requirement and hence maximize its extinction ratio, removing the need for additional halfwave plates and power losses from their imperfections. Following the Pockels for each path, the beams are expanded to a 1/e^2^ diameter of 2.2 mm. A long-pass filter (800-nm cut-off) is mounted upstream of the beam expander to block any undesired residual wavelengths from the ultrafast laser source.

### B. Illumination-scanning optics subsystem

Visible and NIR beams from the multi-laser subsystem are each raised onto 24 inch × 36 inch optical breadboards, one for each twin, by periscopes (Fig. 3). Polarizing beamsplitters are used to both combine the visible and NIR beams into a co-linear beam, and to split the combined beam into two paths (illumination arms 1 and 2). Visible and NIR half-wave plates, each mounted in manual rotation mounts, are used to adjust the laser power (splitting ratio) delivered to illumination arms 1 and 2. Illumination arm 1 is further split into two paths through another half-wave plate and PBS, creating a total of 3 illumination paths. Rotation of the NIR half-wave plates before the polarizing beamsplitter can be used to adjust the relative laser power into illumination arms 1 and 3 (Fig. 3). The path lengths of all 3 illumination arms are routed so that they are equal.

The beams from each illumination arm are sent to 2D scanning galvo mirror positioning systems. The first galvo mirror rapidly scans the beam laterally to synthesize the light-sheet (in the *x*-*y* plane), and the second galvo mirror, which is conjugate to the back pupil of the excitation objective lens, deflects the virtual light-sheet along the (*z*) detection axis. Following the scanning system, each illumination beam passes through a scan lens (achromatic doublet; see Supplementary Note 1); a tube lens; and a low magnification, low NA, long-working-distance excitation objective lens. The distances between pairs of lenses form a 4*f* arrangement (the distance between pairs of lenses are equal to the sum of their focal lengths).

Three excitation objective lenses, mounted orthogonally to each other, direct the illumination light toward the sample (Fig. 3). Depending on the sample properties, any combination of the excitation objectives can be used, either sequentially or simultaneously. Small and/or transparent samples, for example, may benefit from single-sided illumination with 1P excitation; whereas relatively large and thick samples may benefit from the uniform illumination coverage offered by using all three objectives with 2P excitation.

The illumination NA for 1P and 2P is ~ 0.02 and 0.03, respectively, yielding Gaussian-beam light-sheets of average thickness ~ 10 *μ*m across a length of ~ 400 *μ*m, as shown in Fig. 7(d). Scanning of the first galvo yields an effective (*x*-*y*) field-of-view of ~ 400 × 1000 *μ*m^2^. The chosen light-sheet thickness ensures that we are able to resolve single neurons (6 – 8 *μ*m in size) throughout the entire ~ 400 × 800 × 250 (*x-y-z*) *μ*m^3^ brain of zebrafish larva at 5 days-post-fertilization (dpf).^20,28^

Light throughput for each twin, defined as the total measured laser power at the sample from the 3 illumination arms divided by 50% of the measured power at the laser output (for 1:1 split into each twin), for the ultrafast laser (taken at *λ* = 900 nm) is ~ 60%. Light throughput in the visible regime, taken by throughput measurements across each CW laser line is ~ 6%. The lower visible light throughput is expected since most of the illumination-scanning optics were selected to optimize NIR throughput and maximize 2P excitation efficiency (Table SI). Both the ultrafast and CW laser sources are able to simultaneously run experiments on both twins with independent power control.

### C. Detection subsystem

The sheet of fluorescence signal generated at the sample is collected by an orthogonally positioned water-immersion detection objective lens (20×, 1.0 NA), mounted to a piezoelectric (piezo) collar. The high NA objective not only enables high-resolution imaging but also maximizes light-collection efficiency, which is critical for maintaining acceptable signal-to-noise ratio while minimizing the excitation laser power to reduce photodamage to live samples. The fluorescence signal passes through a filter wheel equipped with emission filters to block the excitation light and transmit the fluorescence signal emitted by the sample; the emission filters are optimized for CFP, GFP, mCherry, and Cy5 (see Table SI). A tube lens forms the primary image of the fluorescence signal onto a scientific complementary metal-oxide-semiconductor (sCMOS) camera.

Depending on the sample properties, the detection subsystem can be arranged for upright and/or inverted configurations. For the tests presented here, we have primarily employed an upright configuration, as shown in Fig. 4, that is optimized for imaging neural activity in behaving larval zebrafish (Fig. 5). Switching to an inverted light-sheet detection geometry, or changing the overall magnification is relatively straight-forward, owing to the system’s arrangement of opto-mechanical components (Fig. 2, 3, and 4). The camera is mounted on a rail (Fig. 2 and 4) so that it can be conveniently adjusted for tube lenses of different focal lengths to provide different magnifications.

**FIG. 4.**
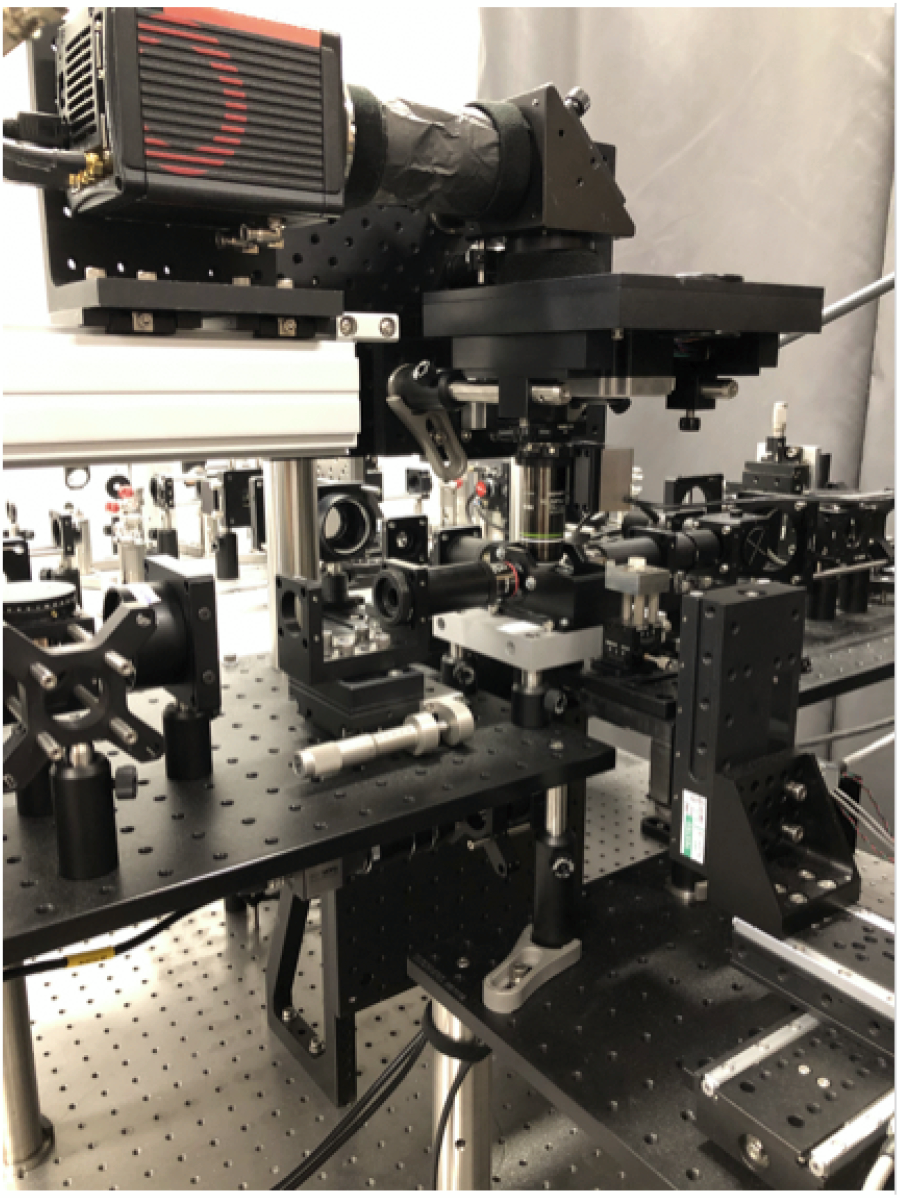
Photograph of an assembled microscope-twin with upright detection.

**FIG. 5.**
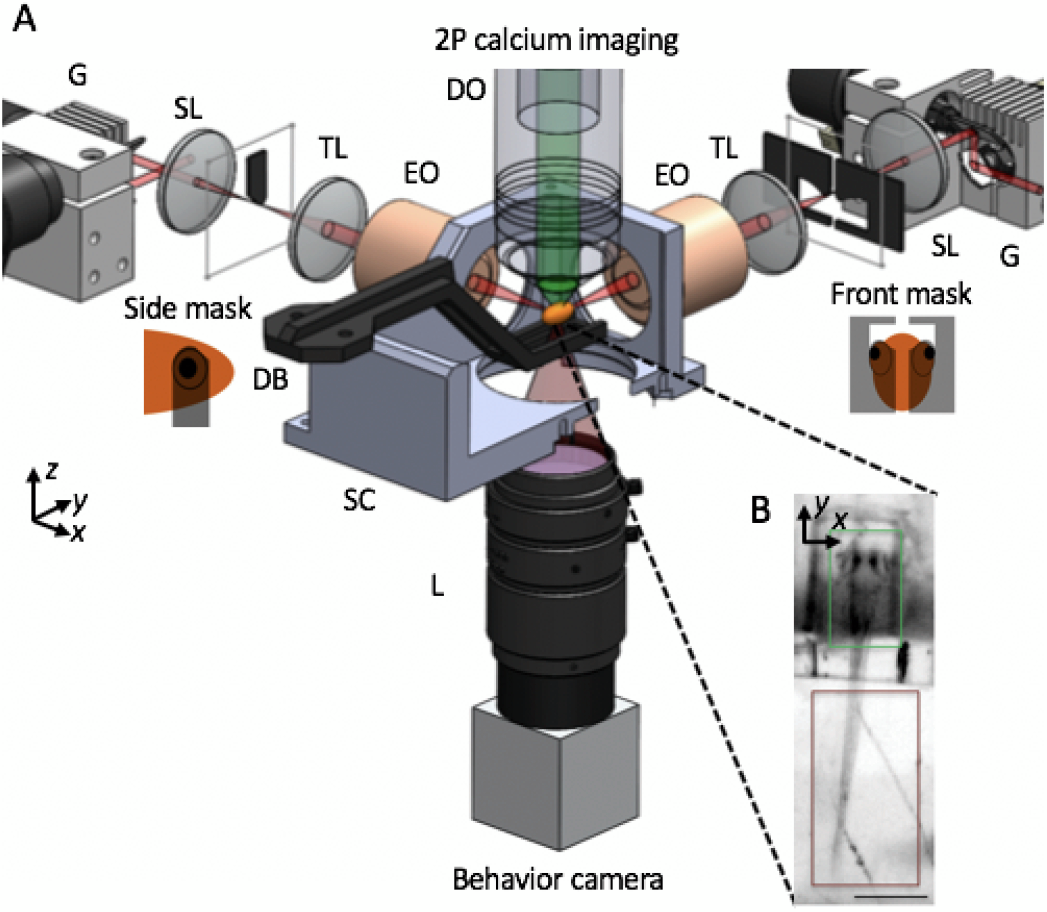
(A) Schematic of apparatus for imaging of neural activity during various behaviors in the larval zebrafish. Sheets of laser light are synthesized by quickly scanning the pulsed-illumination beam (red) with galvo mirrors (G). 2P light-sheets are delivered to the agarose-embedded head of the animal with excitation objectives (EO) from the side and front arms. The side masks cover each eye on the sides of a horizontally oriented zebrafish, while the front mask covers both eyes, enabling access to neurons between the eyes. 2P-excited calcium fluorescence signal is collected through an upright detection objective (DO) and onto a scientific CMOS camera. A triggerable wide-field camera is positioned below the sample chamber (SC) to provide a wide-field, low-resolution view of the sample, as shown in (B). During a typical neural imaging experiment, the zebrafish’s head is immobilized in agarose while the tail is free, thereby permitting the monitoring of zebrafish behavior through tail movement. SL: scan lens; TL: tube lens; DB: dive bar; SC: sample chamber; L: camera lens. The third illumination arm, emission filter, detection TL, scientific camera, light-emitting diode, and filter for behavior channel are not shown. Insets in (B) highlight that the calcium fluorescence channel (green) is recorded from the zebrafish brain, while the behavioral channel (dark red) monitors the tail movement of the animal. Scale bar, (B) 400 *μ*m.

### D. Sample mounting and motion control subsystem

The sample chamber has three side windows for the excitation objectives as well as a bottom window to provide an additional view of the specimen. The sample chamber is open at the top, and is filled with imaging buffer; the open-top allows the detection objective to be liquid-immersed and the sample holder to be inserted. The sample chamber sits on a custom heat exchanger that has circulatory channels for temperature regulated fluid flow, which can be used to keep the medium-filled sample chamber at a specific temperature.

The sample holder is comprised of a caddy that holds the specimen, and a dive bar that holds the caddy. Caddies can be used for agarose embedding of the sample or adapted to specific applications. For example, the caddy for imaging neural activity in behaving zebrafish [Fig. 5(b)] immobilizes the specimen’s head with 1.5% low-melting agarose gel (to record cellular-resolution whole-brain neural activity), leaving the tail free to move (to record swimming behavior). The dive bar is mounted to a dual-axis goniometer, providing rotational motions around the *x*- and *y*-axes; the goniometer is mounted to a motorized 3D stack-up of linear stages (see Table SI), with each stage providing ± 25 mm of travel range. The combination of the two-axis goniometer and 3D stage stack-up allows fine sample positioning so that the illuminated region of interest can be overlapped with the detection objective focal plane.

The flex-SPIM has two different modes of capturing volumetric information from a 3D sample: either by sample-scanning or objective-scanning.

- Sample-scanning mode: the excitation light-sheet and detection objective remain stationary; the sample is moved *via* the *z*-stage of the 3D stage stack-up along the optical axis of the detection subsystem and images are sequentially collected. This approach is the simplest to implement, but its imaging speed is limited by mechanical inertia and communication overhead between the acqusition computer and the z-stage controller. Further, the translational motion of the specimen can compromise normal biology.
- Objective-scanning mode: the movement of the detection objective piezo collar is synchronized with the second galvo mirror of each illumination arm—the light-sheet and piezo collar are scanned axially in concert (with a travel range of ± 500*μ*m set by the piezo collar) and images are sequentially collected (further details are described in next section). This approach enables fast volumetric imaging without moving the specimen, and is pre-ferred for our whole-brain functional imaging and simultaneous behavioral observation studies with zebrafish.

### E. Instrument control module

Each twin is independently controlled with a computer equipped with two Xeon E5-2650 v4 processors and 128 GB of 2400 MHz DDR4 RAM; and seven PCIe slots, enough space for all the control cards. While most of the instrument control and image acquisition routines are done through Micro-Manager,^29^ custom software developed in LabVIEW (National Instruments) is used to independently control each of the 2D scanning galvo mirror systems, and allows precise alignment (size and sweptrate control) of the excitation light-sheet relative to the sample. Collected images are written directly to a dualdisk array consisting of eight 7200 RPM, 4 TB disks.

In the objective-scanning mode, the piezo collar’s controller serves as the master timing source. The analog position-readout of the piezo collar triggers a PicoScope, which is used to generate control signal sequences to synchronize the camera(s) with image capture. The position-readout of the piezo collar is also used to drive the position of the the *z*-galvos. The waveform from the timing output of the scientific CMOS camera controls the AOTF and Pockels, so that the sample is not illuminated during the camera readout. Analog control signals for the galvos and Pockels are appropriately conditioned by individual scaling amplifiers. A schematic of the control signal sequences is shown in (Fig. 6).

**FIG. 6.**
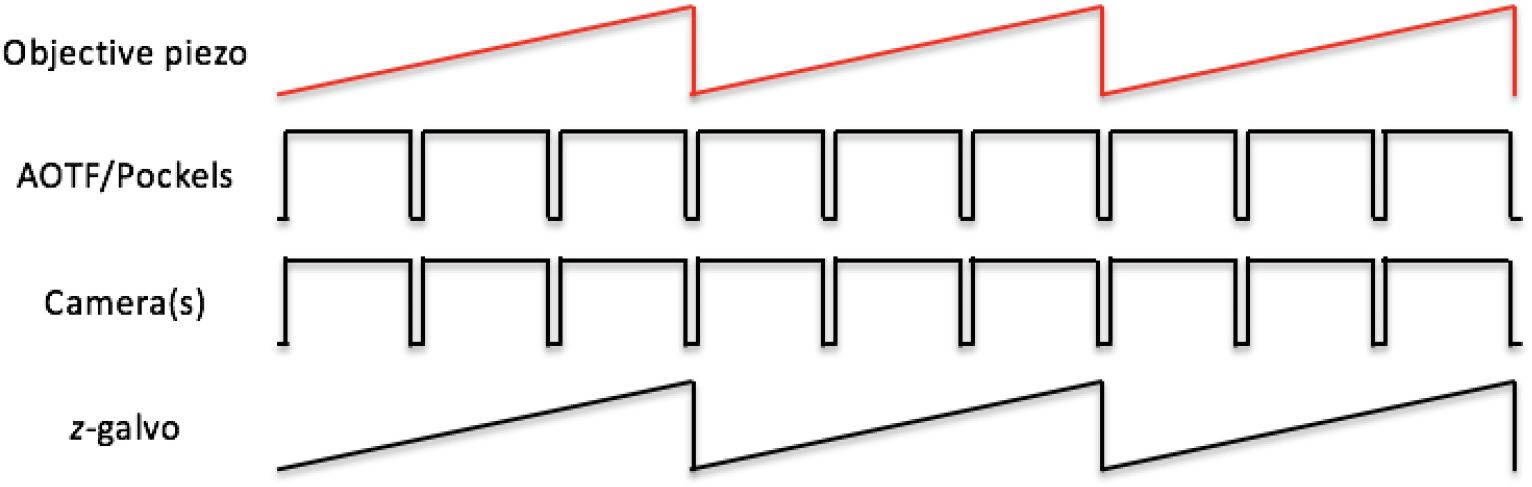
Schematic of control signal sequences for objective-scanning mode. The analog signal representing the position of the objective piezo collar is used as the master timing signal to generate control signals for the imaging cameras (both the scientific CMOS camera and behavior camera). The timing output of the scientific CMOS camera controls the AOTF/Pockels. The number of pulses driving the cameras, shown as 3 in the schematic here, determine the number of individual *z*-plane images to be recorded during a single *z*-scan cycle over the sample. The position signal of the objective piezo collar, appropriately scaled by a scaling amplifier, is also used to drive the *z*-galvo.

### F. Auxiliary module

Illumination can be selectively blocked with masks to avoid photosensitive regions or autofluorescent features in samples. For example, to avoid illuminating the zebrafish eyes while imaging neural activity, the excitation light is physically blocked with a pair of masks on each side of a horizontally positioned zebrafish, and another mask for the front that covers both eyes [Fig. 5(a)]. These masks, fabricated out of black anodized aluminum, are mounted at the image planes of the illumination-scanning optics, each on 2D translational stages to permit their accurate positioning for different specimens, or their complete removal from the illuminated field. A far-red light-emitting diode and a wide-field camera, positioned to view the sample from the bottom, enable view-finding and monitoring the tail behavior during neural imaging of the zebrafish (Fig. 5).

## III. INSTRUMENT PERFORMANCE

We characterized the 3D resolution of the flex-SPIM by measuring the point spread function (PSF) with sub-diffraction fluorescent beads, and then demonstrated the utility of the instrument for investigating biological systems by imaging the beating embryonic zebrafish heart, thick patient-derived tumor organoids, and neural activity in behaving zebrafish. All zebrafish raising and handling procedures followed guidelines established in the Guide for the Care and Use of Laboratory Animals by the University of Southern California, where the protocol was approved by the Institutional Animal Care and Use Committee (IACUC). All zebrafish lines used are available from ZIRC (zebrafish.org).

### A. Resolution characterization

We measured the system PSF with sub-diffraction (175 ± 5 nm diameter) fluorescent beads (PS-Speck Microscope Point Source Kit, P7220, Molecular Probes) embedded in 1.5% agarose, at 44× and 11× magnifications. We computed the averaged PSF from 5 individual bead sub-volumes using Gaussian fits to the intensity profiles. At 44× the averaged lateral and axial full-width-half-maximum (FWHM) ± standard deviation (SD) values are: 1P, 588 nm ± 14 nm and 1.6 *μ*m ± 61 nm, respectively; 2P, 511 nm ± 43 nm and 1.69 *μ*m ± 104 nm, respectively. At 11× the averaged lateral and axial FWHM ± SD values are: 1P, 1.2 *μ*m ± 158 nm and 1.79 *μ*m ± 342 nm, respectively; 2P, 1.23 *μ*m ± 87 nm and 1.65 *μ*m ± 323 nm, respectively. Resolution performance is less than the ideal theoretical diffraction limit associated with the detection optics, mainly stemming from the fact that the experimental spatial sampling, at both magnifications, was not fine enough to meet the Nyquist criterion to exploit the full detection NA.^30^ Despite this practical limitation, the flex-SPIM is still able to achieve submicron, subcellular lateral resolution in both 1P and 2P mode [Fig. 7(c) and 8]. Representative bead images are shown in Fig. 7(a) and (b).

**FIG. 7.**
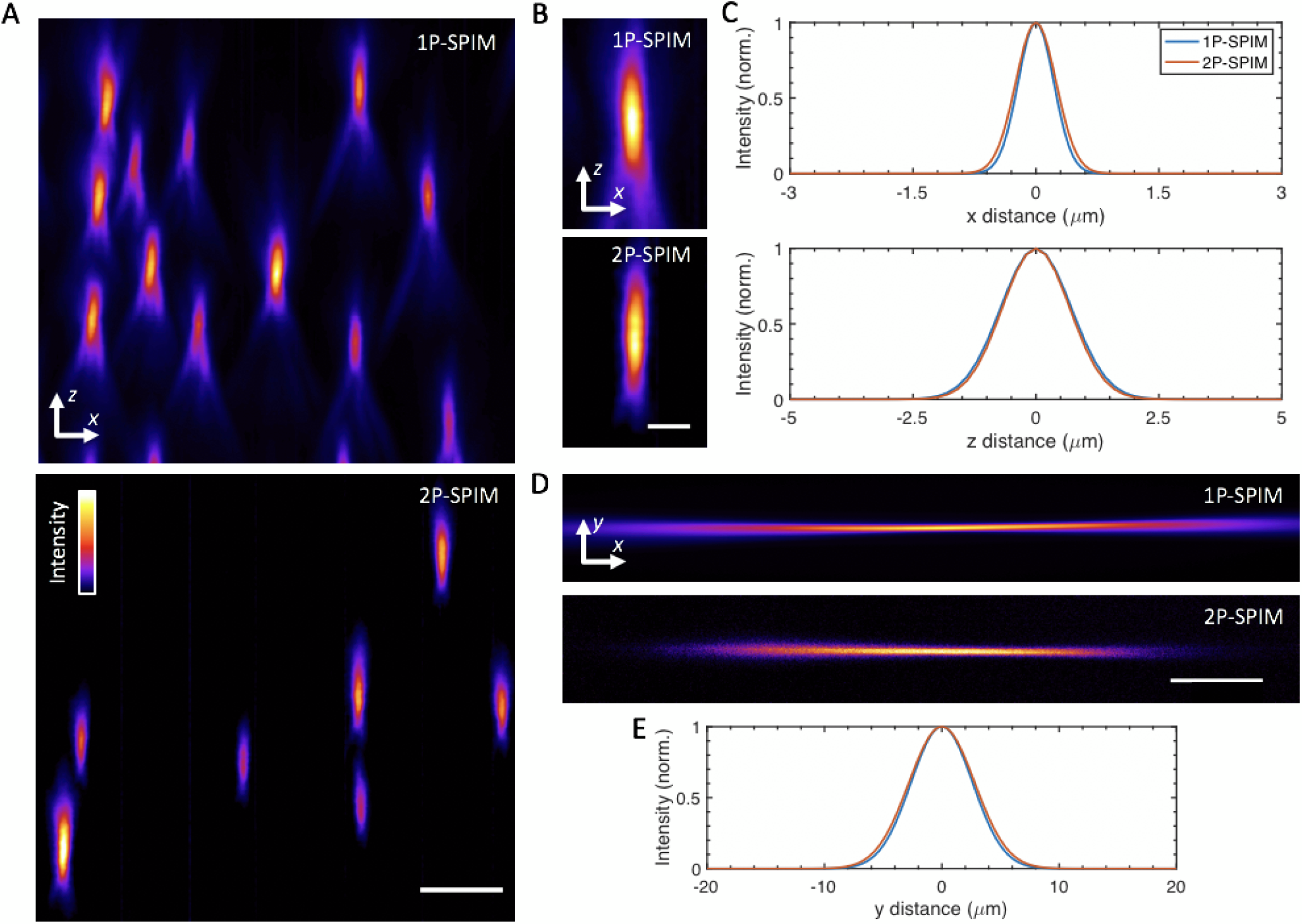
System imaging performance and characterization. (A) *y* maximum-intensity projections of agarose-embedded 175 nm fluorescent beads imaged at 44× magnification, in 1P (top) and 2P excitation mode (bottom). A false-color (fire) look-up table was used to enhance visualization. (B) Selected *y* maximum-intensity projections of sub-diffraction fluorescent beads, in 1P (top) and 2P mode (bottom). (C) Averaged lateral (top) and axial (bottom) full-width at half-maximum (FWHM) extents for the imaged beads, determined by Gaussian fits of 5 bead intensity profiles. The averaged lateral and axial FWHM ± SD values are: 1P, 588 nm ± 14 nm and 1.61 *μ*m ± 61 nm, respectively; 2P, 511 nm ± 43 nm and 1.69 *μ*m ± 104 nm, respectively. (D) Experimental images of fluorescence excited by 1P (top) and 2P (bottom) Gaussian focused beams, which are scanned in the *y* direction to create virtual light-sheets. Images were acquired by illuminating a solution of fluorescein in the sample chamber. (E) Intensity line profiles for the focused beams in (D), taken at the center of focus, with approximate FWHM values: 1P, 6.2 *μ*m; and 2P, 6.6 *μ*m. These FWHM values yield an averaged light-sheet thickness of ~ 10 *μ*m across the 400 *μ*m extent along the *x* direction, centered around the Gaussian focus. Scale bars, (A) 5 *μ*m, (B) 2.5 *μ*m, (D) 150 *μ*m.

**FIG. 8.**
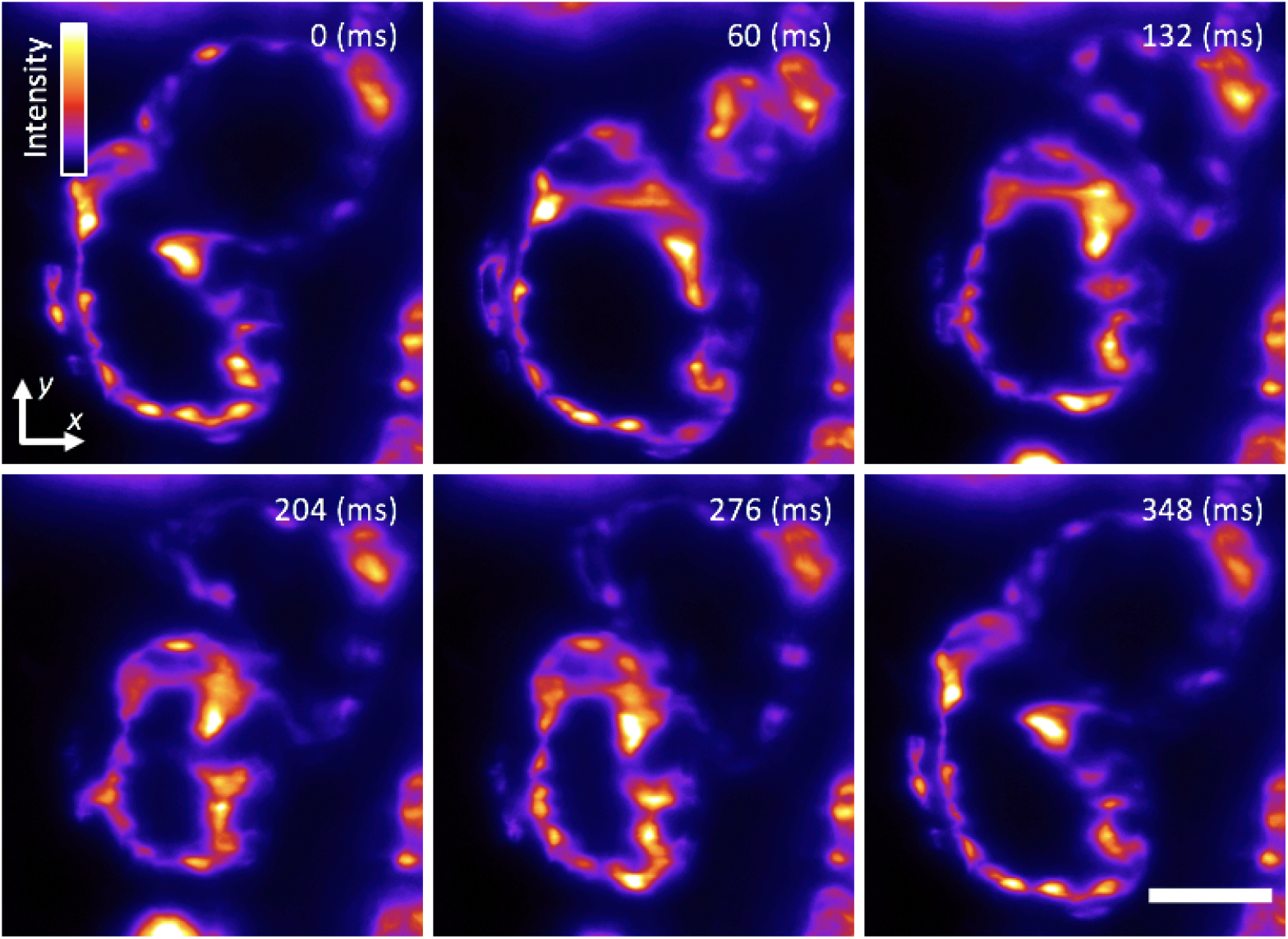
Cardiac light-sheet imaging. Single-plane SPIM recording of the beating heart in a live 5-dpf larval zebrafish with the endocardium fluorescently labeled (GFP), showing 6 distinct time points during the cardiac beating cycle. These subcellular 2D images are comparable to our previous efforts^21^ as well as recent work by others.^31^ A false-color (fire) look-up table was used to enhance visualization. Frames were captured with a magnification of 11× and 5 ms exposure time, at a rate of 85 frames per second. Movie S1 shows a movie of the same data. Scale bar, 50 *μ*m.

### B. Light-sheet imaging of the beating zebrafish heart

The vertebrate heart is a highly dynamic organ that starts to take its form and function early on during development.^32^ To gain insight into how the heart develops, studies of cells in their native dynamic and 3D context in the intact heart are needed. While the zebrafish is an ideal model system because of its optical and genetic accessibility,^33^ imaging is challenged by the heartbeat (2-4 Hz) of over tens of microns in amplitude.^34^ Retroactive synchronization techniques can align the 2D images, by taking advantage of the quasi-periodicity of the heart motion.^21,31,35,36^ We acquired 2D images of the beating heart at high-spatiotemporal resolution in a 5-dpf transgenic larval zebrafish with the vasculature fluorescently labeled. The 1P excitation (*λ* = 488 nm) light-sheet was parked in *z* to optically section through the beating heart of the agarose-embedded sample as we acquired images at 85 frames per second, 11× magnification, and subcellular resolution (Fig. 8 and Movie S1), showing that the flex-SPIM is fully compatible with existing retroactive synchronization techniques.^21,31,35,36^

### C. 3D imaging of thick, patient-derived tumor organoids

3D cell culture systems, such as spheroids or organoids, recapitulate the native physiology of multicellular tissues much better than 2D culture systems.^37^ Multicellular cancer organoids permit the study of disease development and patient-specific response to therapy.^38,39^ Unfortunately, such multicellular systems are scattering and aberrating, making them challenging to image with conventional instruments.

To show the advantages of the flex-SPIM for such opaque and optically heterogenous samples, we imaged, at 11× magnification, chemically fixed, agarose-embedded organoids differentiated from cells derived from a colorectal cancer patient that had been engineered to transgenically express nuclear-localized H2B-GFP [Fig. 9(a) and Movie S2]. 2P-SPIM provides better contrast throughout the imaged volume because the reduced scattering at the longer wavelength (*λ* = 900 nm) enables better-preserved light-sheet shape over longer propagation distances compared to 1P [Fig. 9(b-c); Fig. 9(e)]. Even when the excitation light scatters, the fluorescence signal is still spatially restricted mainly to the central part of the light-sheet (where intensity is the highest) because of the quadratic dependence of the 2P-excited fluorescence signal on the excitation intensity. Thus, by mitigating the scattering-induced thickening of the light-sheet, 2P excitation with the flex-SPIM reveals more clearly-resolved cells than 1P deep in the specimen [Fig. 9(d) and (f)].

**FIG. 9.**
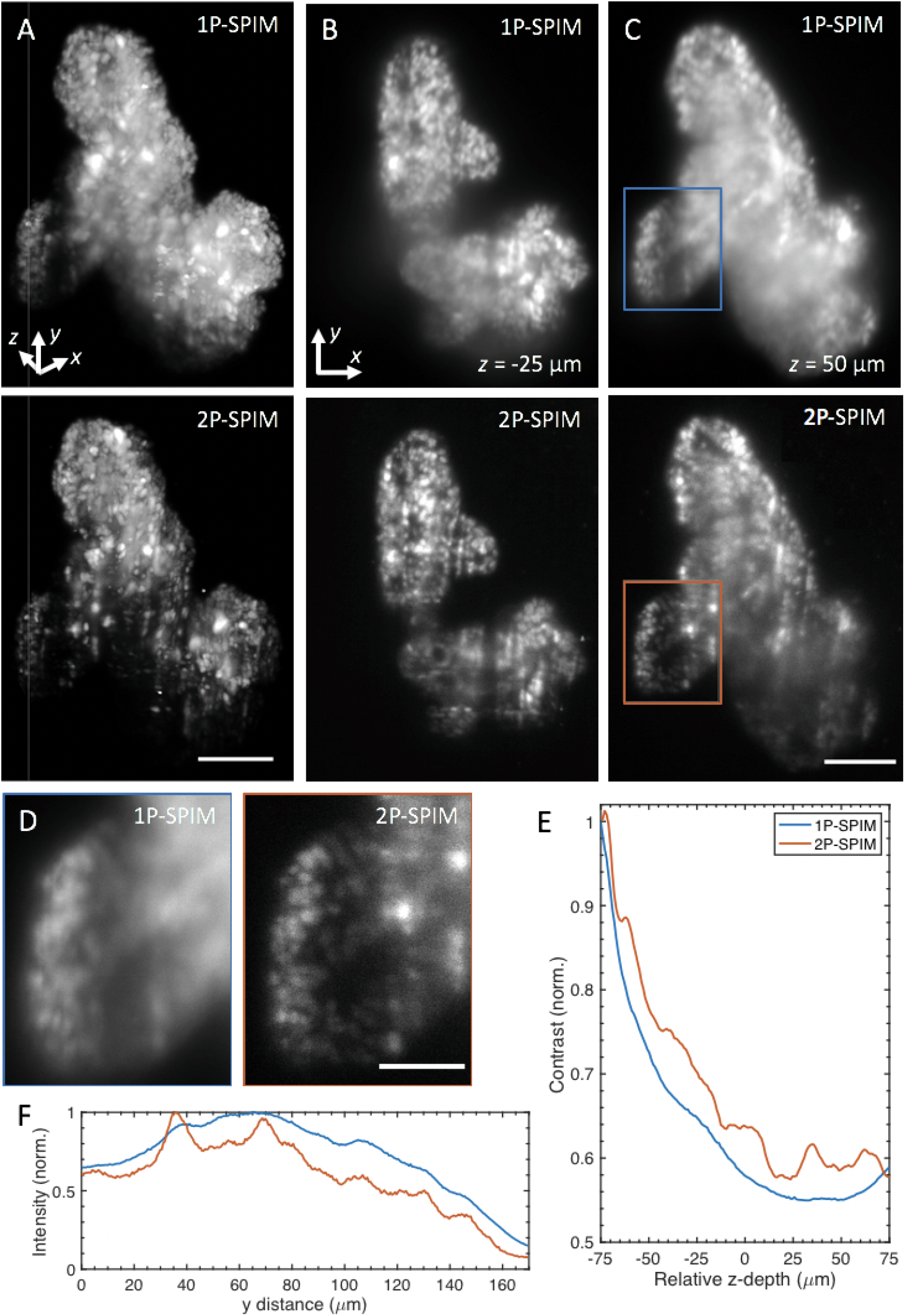
1P- and 2P-SPIM imaging of thick tumor organoids derived from a patient with colorectal cancer. (A) Volume rendering of fixed patient-derived tumor organoids expressing nuclear-localized H2B-GFP recorded in 1P (top) and 2P mode (bottom). Renderings show the reduced background of 2P-SPIM enables better contrast throughout the imaged volume compared to 1P-SPIM. 3D organoid volume of ~ 400 × 550 × 150 (*x-y-z*) *μ*m^3^ captured with a magnification of 11×, 1-*μ*m *z*-steps, and 150 ms exposure time. Movie S2 rotates the 3D-rendered volume of the same datasets. (B) and (C) are *x-y* image slices of (A) at *z* = −25 *μ*m (50 *μ*m from the surface) and *z* = 50 *μ*m (125 *μ*m from the surface), respectively. (D) Magnified images of the boxed regions in (C), for 1P (left) and 2P (right) mode, revealing that 2P-SPIM resolves more cells than 1P-SPIM deep in the sample. (E) Quantification of image contrast as a function of *z*-depth. This plot shows quantitatively the improved contrast of 2P-SPIM over 1P-SPIM throughout the imaged volume in (A). Contrast calculated from the standard deviation of the pixel intensities from each *x-y* image slice and then normalized by the corresponding average image intensity. Each slice (from both modalities) is normalized against the surface slice (*z* = −75 *μ*m) of 1P-SPIM to show the degradation of performance as a function penetration depth. (F) The longer NIR wavelength used in 2P-SPIM minimizes the scattering-induced degradation of the excitation light-sheet over longer propagation distances compared to the visible light used in 1P-SPIM, resulting in better resolution. Plot shows sum intensity along the *x* direction of images in (D) as a function of light-sheet propagation distance *y*. In both intensity profiles, intensity values were normalized by the global maximum. Scale bars, (A, C) 100 *μ*m, (D) 50 *μ*m.

### D. Whole-brain functional imaging of behaving zebrafish

SPIM enables recording whole-brain neural activity in transgenic larval zebrafish.^15,16^ These implementations, however, potentially stimulate the photoreceptors and other photosensitive cells in the retina with the visible excitation wavelengths used during acquisition. Such illumination can reduce visual sensitivity to stimuli and interfere with visually driven processes.^20^ NIR (*λ* = 930 nm) 2P-SPIM overcomes this problem,^20^ achieving a recording depth of 120 *μ*m at a 1 Hz volume rate (sampled by 9 *z*-planes).^17^

We push the depth of 2P light-sheet functional imaging further with the flex-SPIM: more than doubling the volume size while maintaining high spatiotemporal performance (Fig. 10 and Movie S3). By employing a trio of 2P excitation arms with masks to avoid direct laser illumination to the animal’s eyes, we imaged the entire (400 × 800 × 250 *μ*m^3^) brain of a 5-dpf zebrafish expressing a pan-neural calcium indicator (elavl3:H2B-GCaMP6s)^16^ at a 0.5 Hz volume rate (sampled by 52 *z*-planes) with single-cell resolution, and simultaneously monitored swimming behavior with a wide-field camera (Fig. 5). The total average laser power delivered to the sample from the three illumination arms was 490 mW. Imaging was carried out for more than 30 minutes con tinuously, and we observed neither qualitative change in the calcium dynamics nor phenotypic signs of toxicity (Supplementary Note 2).

**FIG. 10.**
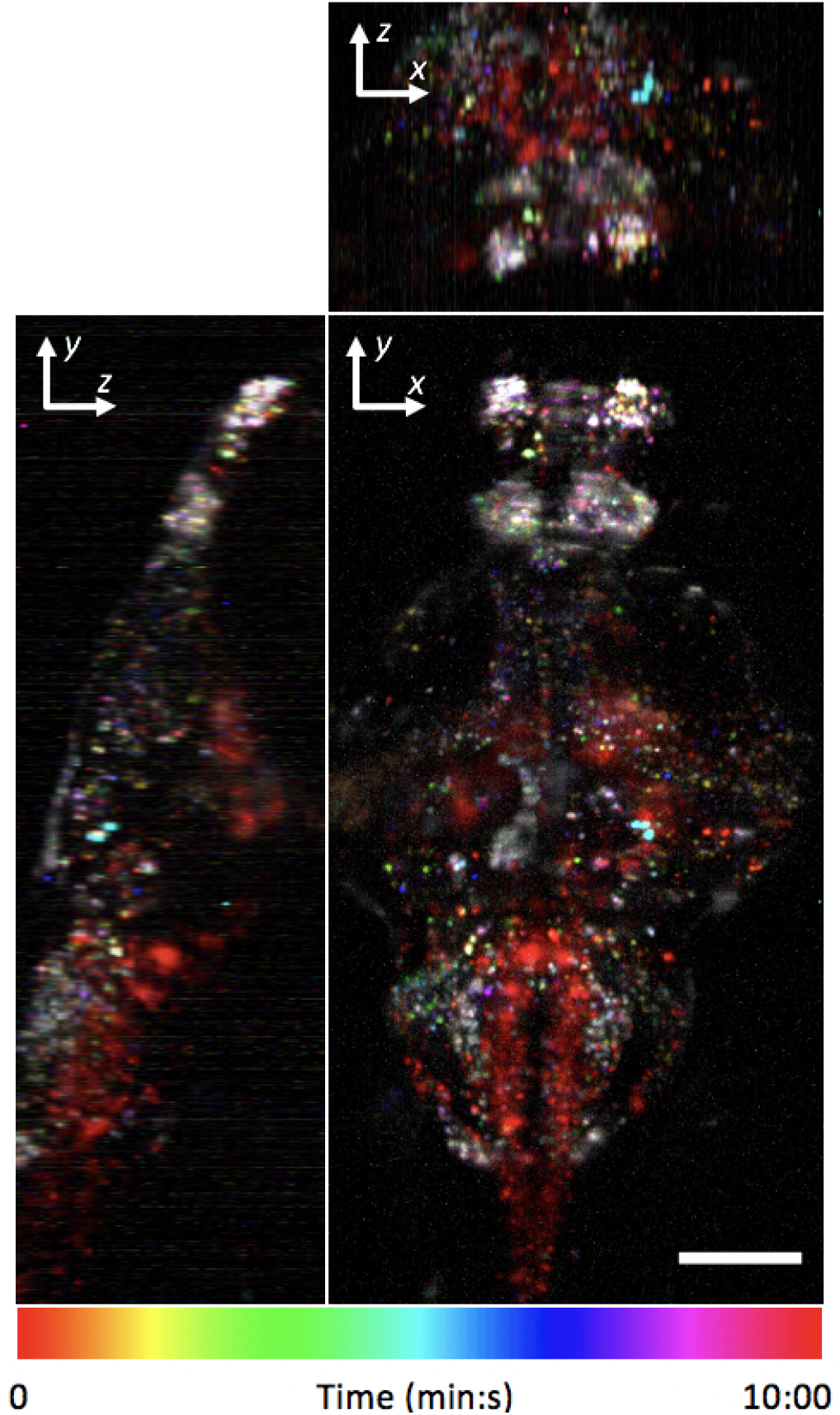
Whole-brain functional imaging at single-cell resolution in behaving 5-dpf transgenic larval zebrafish expressing nuclear-localized calcium indicator elavl3:H2B-GCaMP6s. Maximum-intensity projections of calcium activity are colorcoded in time over the 10-minute recording window. Active neurons that exhibit fluorescence change during the recording appear as colored dots. Volume of 400 × 800 × 250 (*x-y-z*) *μ*m^3^ was sampled by 52 *z*-planes (4.8 *μ*m *z*-steps) at 0.5 Hz with 11× magnification. Movie S3 shows a 3D rendered movie of the same data. Scale bar, 100 *μ*m.

## IV. DISCUSSION

We present the design and construction of an instrument, the flex-SPIM, with two independently controlled light-sheet microscope-twins sharing the same multi-laser source, dramatically cutting the cost of the system. We image a variety of specimens, demonstrating instrument versatility and application-specific customization.

In the same spirit as the OpenSPIM project,^40^ we offer a blueprint for optical developers to build and/or modify the flex-SPIM to serve user needs. A number of further modifications and enhancements could be implemented on the flex-SPIM to further optimize its performance for particular needs. Incoherent structured-illumination from intensity-modulated illumination patterns generated by the AOTF and/or Pockels would enhance contrast in more scattering specimens, but would require additional exposures and post-processing.^41^ Confocal line detection using the rolling shutter of the sCMOS is an efficient alternative to structured-illumination and would allow rejection of nonballistic photons.^42^ Designing a sample chamber rig with both temperature and CO2 control in conjunction with inverted plane-wise detection would allow live organoid imaging. Light-field microscopy could be readily deployed on the flex-SPIM, enabling high-contrast, synchronous volumetric imaging with SPIM-inspired selective volume illumination.^43^

Implementing multispectral imaging on the flex-SPIM would improve signal multiplexing, either on the illumination path by rapid multispectral excitation,^44^ or on the detection path by descanned detection *via* a confocal slit and diffraction grating.^45^ Further improvement is possible with our hyperspectral phasor software (HySP) for unmixing multiple spectrally overlapping fluorophores, even in the face of low signal-to-noise.^46^ The combination of HySP with a multispectral flex-SPIM design could thus enable dynamic visualization and quantitative analysis of many more important components and their interactions in intact specimens at high-resolution over extended durations.

## Appendix: Supplementary material

See supplementary material for a detailed list of the main flex-SPIM parts (Table SI), a panoramic photograph of the flex-SPIM (Figure S1), simulations of scan lens performance (Supplementary Note 1), single-cell neuronal activity extracted from 2P-SPIM whole-brain calcium imaging (Supplementary Note 2), 1P-SPIM imaging of the beating embryonic zebrafish heart (Movie S1), a volume rendering of patient-derived tumor organoids (Movie S2), and a 2P-SPIM recording of neural activity in a behaving animal (Movie S3).

## ACKNOWLEDGMENTS

We are grateful to Sara Madaan for custom LabView software; Matt Jones for electronics help; Peter Luu for zebrafish sample preparations; Seungil Kim and Shannon Mumenthaler (Lawrence J. Ellison Institute for Transformative Medicine of USC) for providing the organoid samples; and Jon Daniels (Applied Scientific Instruments, Inc.) for valuable thoughts on beam-scanning in light-sheet microscopy. Special thanks to Andrey Andreev and the other members of the Translational Imaging Center for insightful discussions, and staff at the Viterbi/Dornsife Machine Shop for technical assistance. This work was supported in part by the National Institutes of Health (1R01MH107238-01), and the Human Frontier Science Program (53-4895-008). K. Keomanee-Dizon was supported in part by the Alfred E. Mann Doctoral Fellowship.

