## Supplementary Material for "A versatile, multi-laser twin-microscope system for light-sheet imaging"

**This PDF includes:**

Table SI

Figure S1

Supplementary Notes 1 and 2

Captions for Movies S1 to S3

**Other Supplementary Materials for this manuscript include the following:**

Movies S1 to S3

#### SUPPLEMENTARY TABLE

**TABLE SI.** List of main flex-SPIM components. Solid model and mechanical drawings for custom components are available upon request.

| Subsystem/ module | Component | Catalogue number | Vendor | Notes |
|---|---|---|---|---|
| | 5 foot × 10 foot, anti-vibration optical table | INTEGRITY 4 VCS 510-8 | Newport | |
| CW Laser bank | OBIS 445 nm LX | 1185051 | Coherent | Power: 75 mW |
| | OBIS 488 nm LX | 1220123 | Coherent | Power: 150 mW |
| | OBIS 561 nm LS | 1280720 | Coherent | Power: 150 mW |
| | OBIS 647 nm LX | 1196627 | Coherent | Power: 120 mW |
| | $f$ = 20 mm, 12.5 mm diameter lens | 47-661 | Edmund Optics | Beam expansion |
| | $f$ = 8 mm, 12.24 mm diameter lens | C240TME-A | Thorlabs | |
| | 25 mm diameter broadband mirror | 87-371 | Edmund Optics | Beam combining |
| | Di02-R561 dichroic beamsplitter | Di02-R561-25x36 | Semrock | Beam combining |
| | LM01-503-25 LaserMUX dichroic beamsplitter | LM01-503-25 | Semrock | Beam combining |
| | LM01-466-25 LaserMUX dichroic beamsplitter | LM01-466-25 | Semrock | Beam combining |
| | 1 inch diameter broadband dielectric mirrors | BB1-E02 | Thorlabs | Broadband (400-750 nm) dielectric beam steering mirror |
| | 30 mm cage cube-mounted variable beamsplitter | VA5-PBS251 | Thorlabs | Visible (420-680 nm) polarization optics |
| | 0.5 inch diameter mounted achromatic half-wave plate | AHWP05M-600 | Thorlabs | Visible (400-800 nm) polarization optic |
| | AOTF | AOTFnC-400.650-TN | AA Quanta Tech, Optoelectronic | Tellurium dioxide crystal<br>Number of channels: 8<br>Wavelength range: 400-650 nm<br>Transmission: > 90%<br>Aperture: 3 $mm^2$<br>Spectral resolution: 1-4 nm<br>Tuning time: < 4 µs |
| | Driver for AOTF | MPDS8C-D66-22-74.158 | AA Quanta Tech | Number of channels: 8<br>Communication: USB, RS232, RC03<br>Extinction ratio: 120 dB |
| | RF cable SMA connectors | CBL-SAM200SAM-RG223 | AA Quanta Tech | |
| | STAB, cable, SMC connectors | CBL-SCF200SCF-RG316 | AA Quanta Tech | |

| Subsystem/ module | Component | Catalogue number | Vendor | Notes |
|---|---|---|---|---|
| NIR femtosecond-pulsed laser | Insight DS+ | 90044047 | Spectra Physics | Tuning range: 680-1300 nm<br>Tuning. range: 680-1300 nm<br>Repetition rate: 80 ± 0.5 MHz |
| | 30 mm cage cube-mounted variable beamsplitter | VA5-PBS252 | Thorlabs | NIR (690-1000 nm) polarization optics |
| | 1 inch diameter broadband dielectric mirrors | BB1-E03 | Thorlabs | Broadband (750-1100 nm) dielectric beam steering mirror |
| | E-O Modulator 2.7mm Aperture | 350-80-02 KD*P | Conoptics | |
| | Galilean beam expander | BE02-05-B | Thorlabs | 2×-5× optical beam expander |
| | 800 nm long-pass filter, 1 inch diameter | FEL0800 | Thorlabs | |
| Illumination-scanning optics | 24 inch × 36 inch optical breadboard | B2436F | Thorlabs | |
| | 66 mm construction rail | XT66-500 | Thorlabs | Periscope |
| | 1 inch kinematic mirror mount | KM100 | Thorlabs | |
| | 45° elliptical mirror mount | H45E1 | Thorlabs | |
| | VIS elliptical mirror | BBE1-E02 | Thorlabs | |
| | NIR elliptical mirror | BBE1-E03 | Thorlabs | |
| | 1 inch pedestal post, 1inch long | RS1P8E | Thorlabs | |
| | clamping platform for 66 mm rail | XT66C4 | Thorlabs | |
| | 0.5 inch diameter silver mirror | PF05-03-P01 | Thorlabs | Beam steering mirror |
| | 0.5 inch diameter mounted achromatic half-wave plate | AHWP05M-600 | Thorlabs | Visible (400-800 nm) polarization optic |
| | 30 mm cage cube-mounted variable beamsplitter | VA5-PBS252 | Thorlabs | NIR (690-1000 nm) polarization optics |
| | 2D scanning galvo mirror positioning system | GVSM002 | Thorlabs | |
| | Mounting adapter for 2D galvo system | GCM102 | Thorlabs | |
| | $f$ = 150 mm, 50 mm diameter lens | VIS-NIR 49-391-INK | Edmund Optics | Scan lens (achromatic doublet) |
| | $f$ = 200 mm, 50 mm diameter lens | VIS-NIR 49-392-INK | Edmund Optics | Tube lens |
| | 5×, 0.10 NA, 23 mm WD objective lens | LMPLN5XIR LWD M PLAN | Olympus | Excitation objective lens |

| Subsystem/ module | Component | Catalogue number | Vendor | Notes |
|---|---|---|---|---|
| Detection | 20×, 1.0 NA, 2 mm WD objective lens | XLUMPLFLN-W | Olympus | Detection objective lens |
| | Filter wheel | Lambda 10-B | Sutter Instrument | Twin 1: 32 mm diameter<br>Twin 2: 25 mm diameter |
| | Filter wheel controller | Lambda 10-3 | Sutter Instrument | |
| | 609/54 nm BrightLine single-band bandpass filter | FF01-609/54-32 | Semrock | Emission filter set for Twin 1 |
| | 680/42 nm BrightLine single-band bandpass filter | FF01-680/42-32 | Semrock | |
| | 525/50 nm BrightLine single-band bandpass filter | FF03-525/50-32 | Semrock | |
| | 470/28 nm BrightLine single-band bandpass | FF01-470/28-32 | Semrock | |
| | 609/54 nm BrightLine single-band bandpass filter | FF01-609/54-25-STR | Semrock | Emission filter set for Twin 2 |
| | 680/42 nm BrightLine single-band bandpass filter | FF01-680/42-25-STR | Semrock | |
| | 525/45 nm single-band bandpass filter | FF01-525/45-25-STR | Semrock | |
| | 472/30 nm BrightLine single-band bandpass filter | FF02-472/30-25-STR | Semrock | |
| | $f$ = 100 mm, 50 mm diameter lens | VIS-NIR 49-284-INK | Edmund Optics | Tube lens (for 11× magnification) |
| | $f$ = 400 mm, 75 mm diameter lens | VIS-NIR 88-598-INK | Edmund Optics | Tube lens (for 44× magnification) |
| | ORCA-Flash4.0 V3 Digital CMOS camera | C13440-20CU | Hamamatsu | Twin 1<br>Number of pixels: 2048 × 2048<br>Pixel size: 6.5 μm2<br>Full resolution frame rate:<br>100 frames/s (Camera link)<br>40 frames/s (USB)<br>Quantum efficiency: 82% peak |
| | ORCA-Flash4.0 V2 Digital CMOS camera | C11440-22CU | Hamamatsu | Twin 2<br>Similar specifications to camera on Twin 1 |
| Sample mounting | Sample chamber | Custom | Protolabs | Material: Delrin 150 black acetal homopolymer |

| Subsystem/ module | Component | Catalogue number | Vendor | Notes |
|---|---|---|---|---|
| | Clamping rings | Custom | Protolabs | Material: Delrin 150 black acetal homopolymer; used to clamp glass windows to sample chamber |
| | Heat exchanger | Custom | Protolabs | Material: Aluminum 6061-T651 or Cooper |
| | Caddy | Custom | Protolabs | Material: WaterShed XC 11122 |
| | Dive bar | Custom | Protolabs | Material: Stainless steel 316/316L |
| | Dive bar to goniometer mount | Custom | Protolabs | Material: Stainless Steel 316/316L |
| | Dual-axis goniometer | GN2/M | Thorlabs | |
| | Metric baseplate | UBP2/M | Thorlabs | |
| | Breadboard | MB1012 | Thorlabs | For sample stack-up |
| | High-purity silicone Rubber .010" thick | 87315K62 | McMaster-Carr | Material: 55A Durometer; gasket used to clamp tightly seal glass windows to sample chamber |
| | High-purity silicone Rubber .020" thick | 87315K63 | McMaster-Carr | |
| | 40mm glass coverslips | 10200-060 | VWR | Used as bottom window for sample chamber |
| | 31mm glass coverslips | NC1491415 | Fisher Scientific | Used as side windows for sample chamber |
| Motion control | nPFocus1000 piezo stage | 3715250 | nPoint | 1000 μm travel |
| | Controller LC.400 | 200761 | nPoint | 1 Axis |
| | LS-50 linear stage; 16 TPI (z-stage) | LS-50-AMERL | Applied Scientific Instrumentation | LS-50 3D stage stack-up |
| | Linear encoder option for z-drive (to attaining resolutions down to 50 nm) | LE-Z | Applied Scientific Instrumentation | |
| | LS-50 linear stage; 4 TPI (x- and y-stages) | LS-50-BMERL | Applied Scientific Instrumentation | |
| | Mount/bracket for linear stages | LS-5013 | Applied Scientific Instrumentation | |
| | Plate for attaching linear stage to breadboard | LS-5012 | Applied Scientific Instrumentation | |
| | Tiger controller, with xyz cards and joysticks | TG16_BASIC | Applied Scientific Instrumentation | |
| Instrument control | Supermicro motherboard with Intel C612 AHCI SATA controller | X10DRH-CT | CDW-G | See (1) below for specifications |
| | NI PXIe-1073 Integrated MXIe, 5 peripheral slots, PCIe-8361, 3 m cable | 781161-01 | National Instruments | |

| Subsystem/ module | Component | Catalogue number | Vendor | Notes |
|---|---|---|---|---|
| | Power cord, AC, U.S., 120 VAC, 2.3 m | 763000-01 | National Instruments | |
| | NI PXIe-6363, X Series DAQ (32 AI, 48 DIO, 4 AO) | 781056-01 | National Instruments | |
| | CB-68LPR I/O connector block | 777145-02 | National Instruments | |
| | SHC68-68-EPM shielded cable, 68-D-type to 68 VHDCI offset, 2 m | 192061-02 | National Instruments | |
| | Mainframe with RS-232 computer interface | SIM900 | Stanford Research | |
| | Scaling amplifier | SIM983 | Stanford Research | |
| | PicoScope 2000 Series oscilloscope | 2207B | Pico Technologies | 2 channel; 70MHz |
| | BNC breakout box | PR35B32CMB | L-com | |
| Auxiliary | Monochrome CMOS camera | DCC3240M | Thorlabs | Behavior camera |
| | USB 3.0 I/O Cable | CAB-DCU-T3 | Thorlabs | Cable for triggering |
| | 35-50 mm fixed focal length camera lens | MVL50M23 | Thorlabs | |
| | T-Cube LED Driver with Trigger Mode | LEDD1B | Thorlabs | Far-red LED |
| | 780 nm, 200 mW mounted LED, 800 mA | M780L3 | Thorlabs | |
| | 1” diameter longpass filter | FEL0750 | Thorlabs | Cut-on wavelength: 750 nm |

(1): Windows 7 Professional w/SP1 64-Bit
1× SMC SuperChassis
CSE-836BE1C-R1K03B
2× Intel Xeon E5-2650 V4 CPUs, 2.2GHz, 12-Core, 30M Cache, 105W
4× 32GB DDR4 2400MHz ECC Registered DIMMs (128GB Installed)
2× 10GbE NIC Ports - Intel X540 dual-port LAN, RJ45 (onboard)
1× Integrated IPMI 2.0 with dedicated LAN
1× MSI 2GB GDDR5 64-Bit
GPU/Videocard, N730K-2GD5LP/OC
8× Empty 3.5"" Drive Bays with trays
1× H/W RAID Controller, LSI 3108, 2GB DDR3 Cache (S3108L-H8IR-16DD onboard)
Supported RAID Levels 0, 1, 5, 6, 10, 50, 60
2× 1000W Redundant Hot-Swap Power
1× Hot-Swap 512GB SATA 6Gb/s OS SSD drive, 2.5"", Samsung 850 Pro
8× Hot-Swap 4TB SATA 6Gb/s data drives, 7200 RPM, Seagate ST4000NM0035
1× DVDRW Slim Black SATA Samsung, SN-208FB/BEBE LG - #GTC0N

**SUPPLEMENTARY FIGURES**

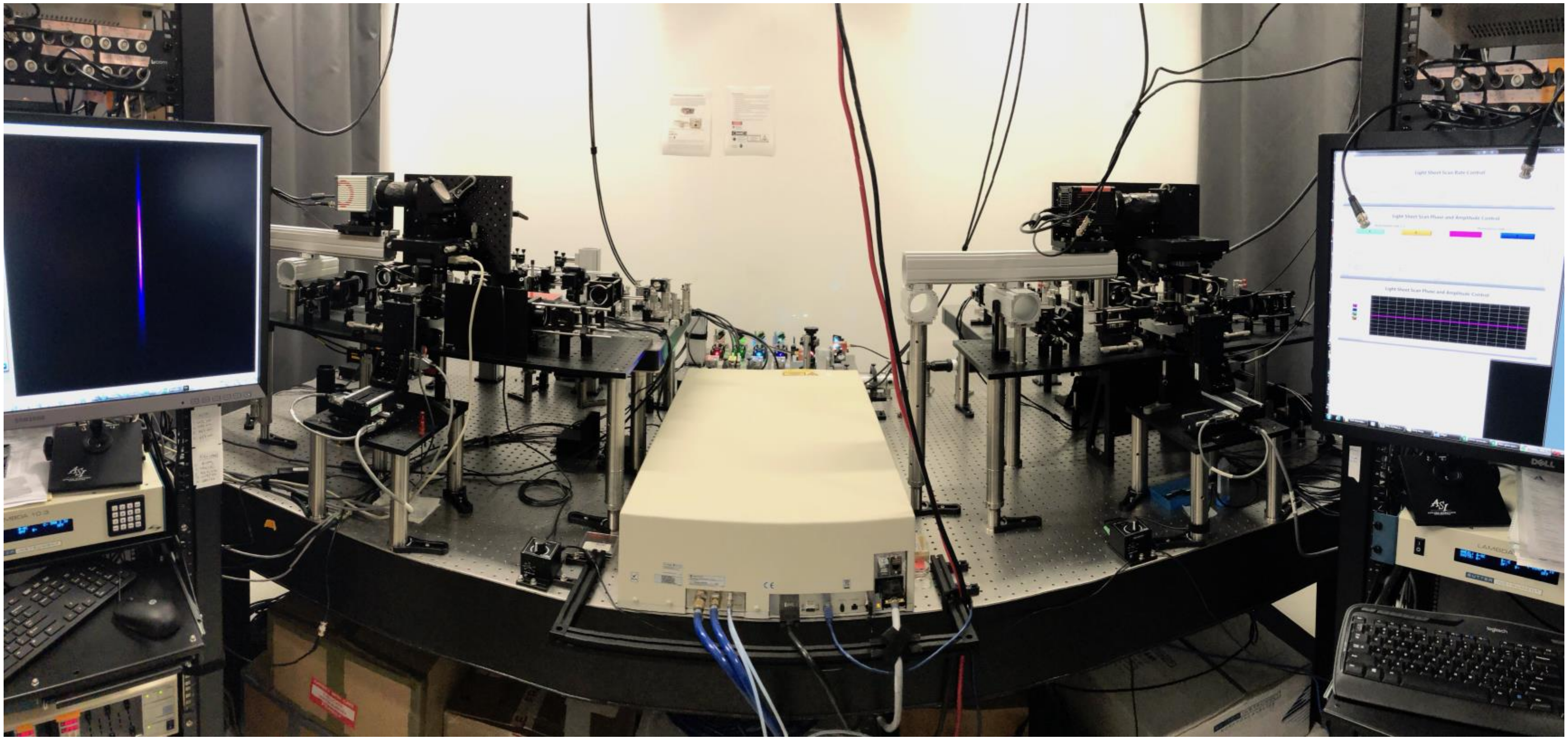

**FIG. S1.** Panoramic photograph of the assembled twin-microscope system, showing the multi-laser subsystem shared between microscope-twin-1 (right) and microscope-twin-2 (left). The corresponding electronic and instrument control racks are shown beside each microscope-twin.

### SUPPLEMENTARY NOTES

#### Supplementary Note 1

##### Simulations of scan lens performance using an achromatic doublet

We chose an achromatic doublet lens as a scan lens, instead of using specially-designed scan lenses which are more expensive. This decision was based on computer-aided optical modeling (Zemax, Radiant Zemax LLC) of commercially available lenses, showing that the achromatic doublet lenses have sufficient performance for our needs. We considered $1/e^2$ Gaussian beam diameters ≥ 1.5 mm and ≤ 4 mm; excitation wavelengths corresponding to 445 nm, 448 nm, 561 nm, 647 nm, 910 nm, and 1040 nm; and a maximum scan angle of ± 4°, which is equivalent to sweeping out a 2.56 $mm^2$ illuminated plane at the sample (given the illumination tube lens and excitation objective described above)—far exceeding the desired ~ 0.5 × 1 $mm^2$ illuminated field. Being ultra-conservative about the maximum field angle ensured minimal aberrations as well as the flexibility to illuminate even larger areas (> 0.5 × 1 $mm^2$), if desired. The chosen achromatic doublet does a surprisingly good job of broadband (445-1040 nm) diffraction-limited performance over a large field angle, as illustrated in Fig. S2. Further optimization could be achieved with the use of custom[47] or more sophisticated lens design,[48,49] such as the Plössl (a pair of symmetric achromatic doublets opposing each other);[48] however, the cost would scale proportionally with each scan lens.

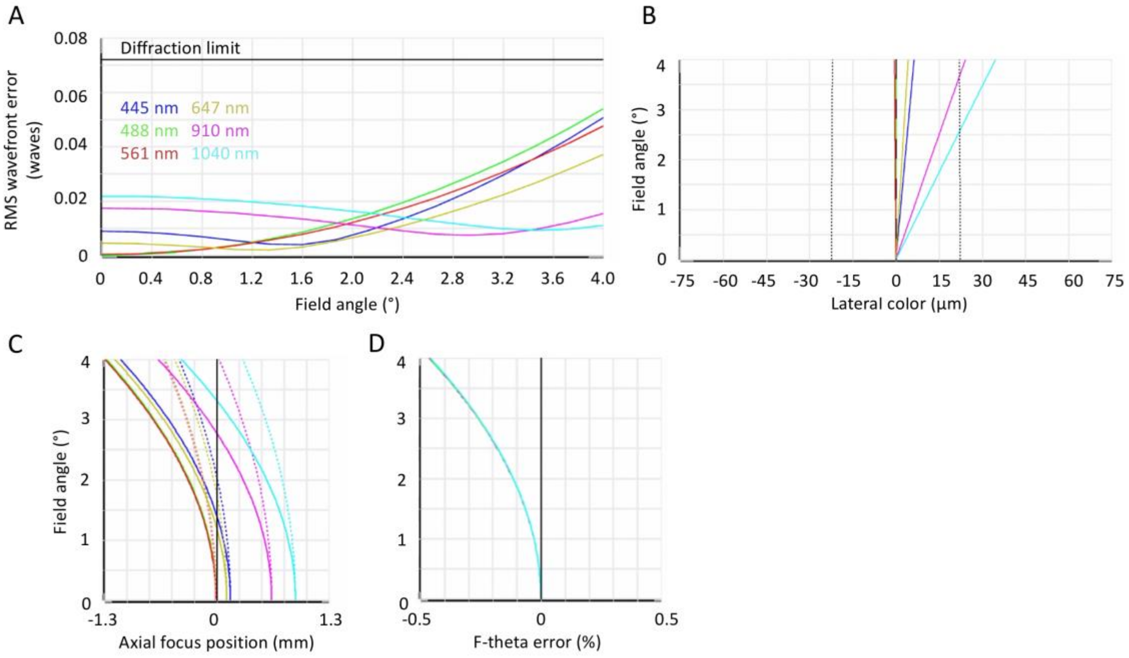


**FIG. S2.** Simulations of the performance of the scan lens used, achromatic doublet ($f$ = 150 mm, 50 mm diameter, VIS-NIR 49-391, Edmund Optics) using a $1/e^2$ beam diameter of 4 mm. (A) Transmitted wavefronts [measured by root mean squared (RMS) wavefront error] remain diffraction-limited, well beyond the desired scan range (for reference, ± 2° is equivalent to a 1.25 $mm^2$ illuminated field at the sample). (B) Using a 1000 nm Airy disc as reference, lateral color remains diffraction-limited for the desired ± 2° scan range; for scan angles > 2.5°, chromatic aberrations occur at 910 nm and 1040 nm. (C) Field curvature for the tangential and sagittal beam components, respectively represented by the solid and dashed lines. At the maximum scan angle of ± 4°, the field curvature is 1.3 mm for the tangential and 0.52 mm for the sagittal components; at the sample, this curvature translates to ~ 0.9 µm for the tangential and ~ 0.4 µm for the sagittal components, both of which are minor deviations relative to the illuminated field. (D) F-theta distortions are minimal: < 0.5% at the largest scan angle.

#### Supplementary Note 2

##### Single-cell neuronal activity extracted from 2P-SPIM whole-brain calcium imaging

We used a total of 490 mW laser power (from 3 illumination arms) at $\lambda$ = 930 nm to image a 5-dpf transgenic larval zebrafish (presented in Fig. 10) continuously for 30 minutes. Neurons were manually selected from the 3D time-lapse dataset [Fig. S3(a)]. Then, the mean fluorescent intensity for each selected neuron was used to calculate the relative fluorescence variation ($\Delta F/F$) over the 30-minute time window, where F is the baseline fluorescence and $\Delta F$ is the fluorescence change due to neural activity. $\Delta F/F$ time traces from the neurons in Fig. S3 show no qualitative change to the calcium dynamics over the 30-minute window, as well as no decay in peak intensities (from activity) as a function of time. Assessment of the 4D dataset showed no structural changes to neurons within the brain [Fig. S3(a)]. In addition, assessment of the low-resolution, wide-field tail channel showed no detrimental behavioral changes (data not shown). These observations suggest that no functional effect of phototoxicity on neural activity was observed from the imaging conditions used.

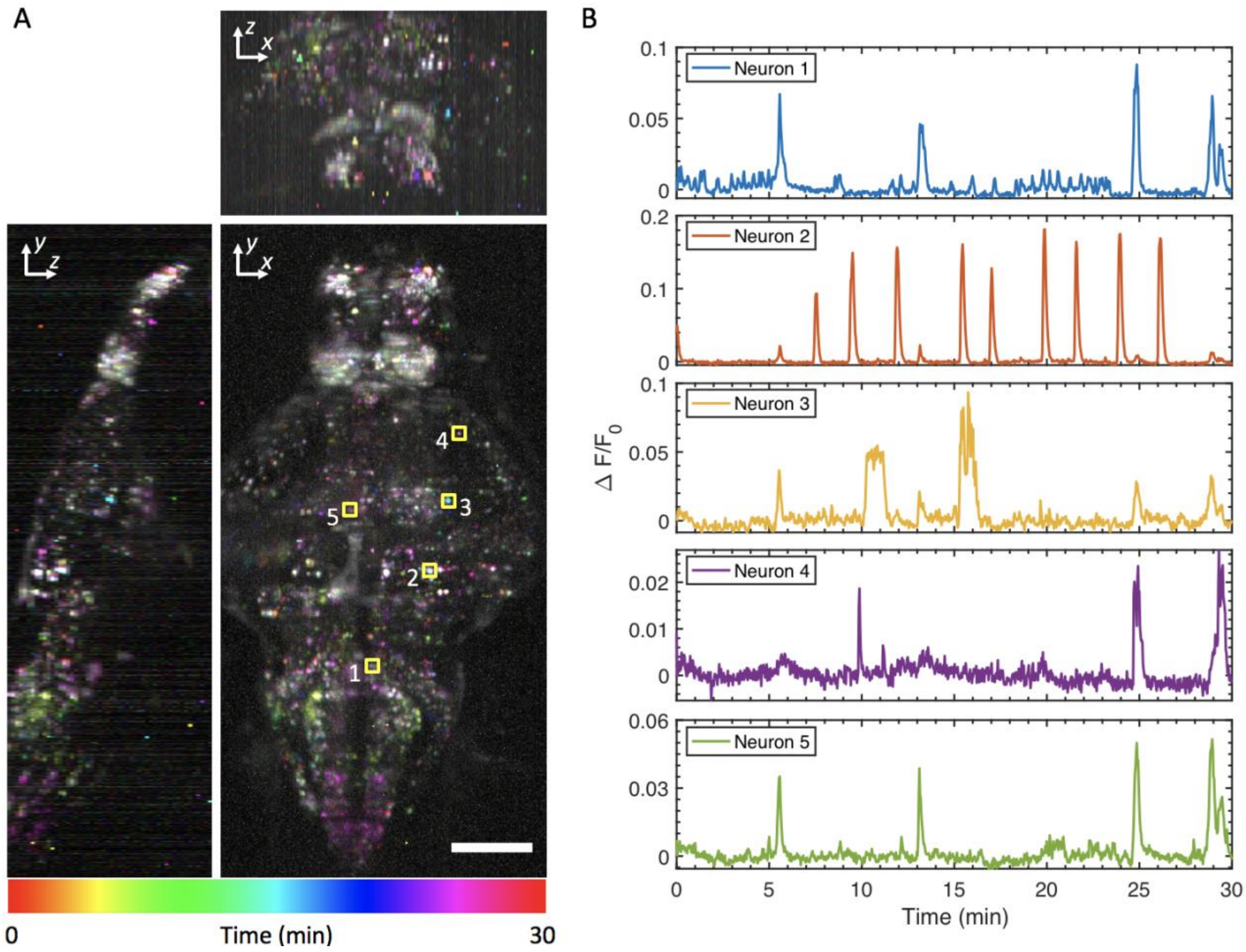


**FIG. S3.** (A) Maximum-intensity projections of calcium activity are color-coded in time over the 30-minute recording window. Same animal and volumetric acquisition parameters as in Fig. 10, showing no phenotypic signs of phototoxicity from illumination at 490 mW laser power. (B) $\Delta F/F$ time traces from the manually selected neurons in (A), showing no qualitative difference in the calcium signal dynamics throughout the 30-minute recording window. Scale bar, (A) 100 µm.

**MOVIE CAPTIONS:**

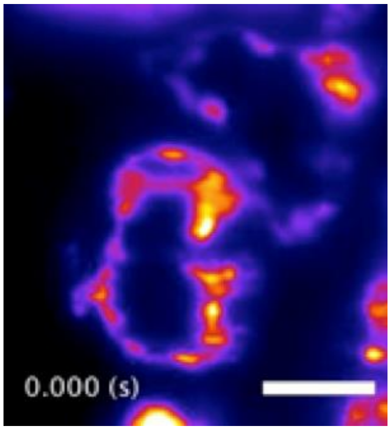


**Movie S1.** Light-sheet imaging of the dynamic motion of the beating heart of a 5-dpf transgenic larval zebrafish. Same dataset as presented in Fig. 8. Frames were captured at 85 Hz. Scale bar, 50 μm.

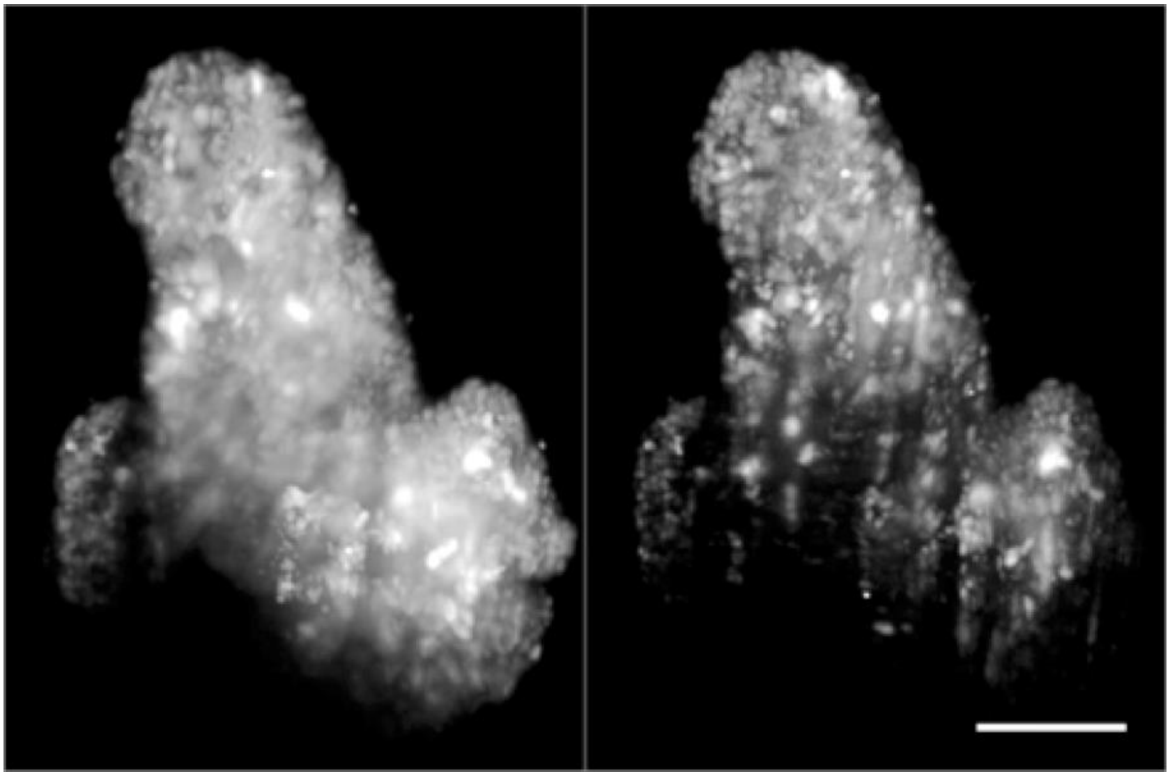

**Movie S2.** Volume rendering of fixed patient-derived tumor organoids expressing H2B-GFP, comparing images taken with one-photon (left) and two-photon-excitation SPIM (right). Volumes are rotated around the *y* and *x* axes. Same datasets as presented in Fig. 9. Scale bar, 100 μm.

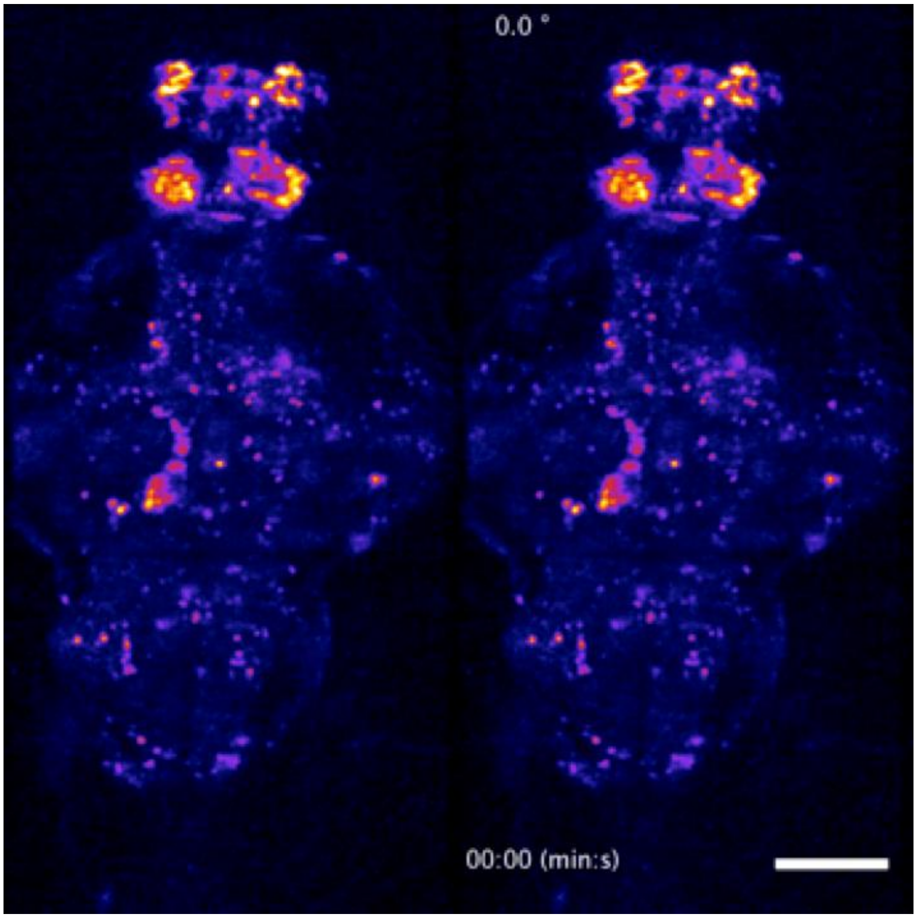


**Movie S3.** Dorsoventral (left) and rotating (right) maximum-intensity projections of a time-lapse recording of the whole-brain of the a 5-dpf transgenic larval zebrafish. Same dataset as presented in Fig. 10. Two-photon whole-brain functional light-sheet imaging was performed at a volumetric rate of 0.5 Hz. The video loops a 5-minute recording of large-scale neural activity in the behaving animal. Scale bar, 100 μm.
